# Interactions between fungal hyaluronic acid and host CD44 promote internalization by recruiting host autophagy proteins to forming phagosomes

**DOI:** 10.1101/2020.04.20.047621

**Authors:** Sheng Li Ding, Aseem Pandey, Xuehuan Feng, Jing Yang, Luciana Fachini da Costa, Roula Mouneimne, Allison Rice-Ficht, Samantha L. Bell, Robert O. Watson, Kristin Patrick, Qing-Ming Qin, Thomas A. Ficht, Paul de Figueiredo

**Author notes:** These authors contributed equally (Sheng Li Ding, Aseem Pandey, Xuehuan Feng). Correspondence (Q.M.Q.), (T.A.F.), (P.d.F.).

## Abstract

Phagocytosis and autophagy play critical roles in immune defense. *Cryptococcus neoformans* (Cn), a fungal pathogen that causes fatal infection, subverts the host autophagy initiation complex (AIC) and its upstream regulatory proteins, to promote its phagocytosis and intracellular parasitism of host cells. The mechanisms by which the pathogen engages host AIC proteins remain obscure. Here, we show that the recruitment of host AIC proteins to forming phagosomes is dependent upon the activity of CD44, a host cell surface receptor that engages fungal hyaluronic acid (HA). This interaction elevates intracellular Ca^2+^ concentrations and activates CaMKKβ and its downstream target AMPKα, which results in activation of ULK1 and the recruitment of AIC components. Moreover, we demonstrate that HA-coated beads efficiently recruit AIC components to phagosomes. Taken together, these findings show that fungal HA plays a critical role in directing the internalization and productive intracellular membrane trafficking of a fungal pathogen of global importance.

**Graphical Abstract:** 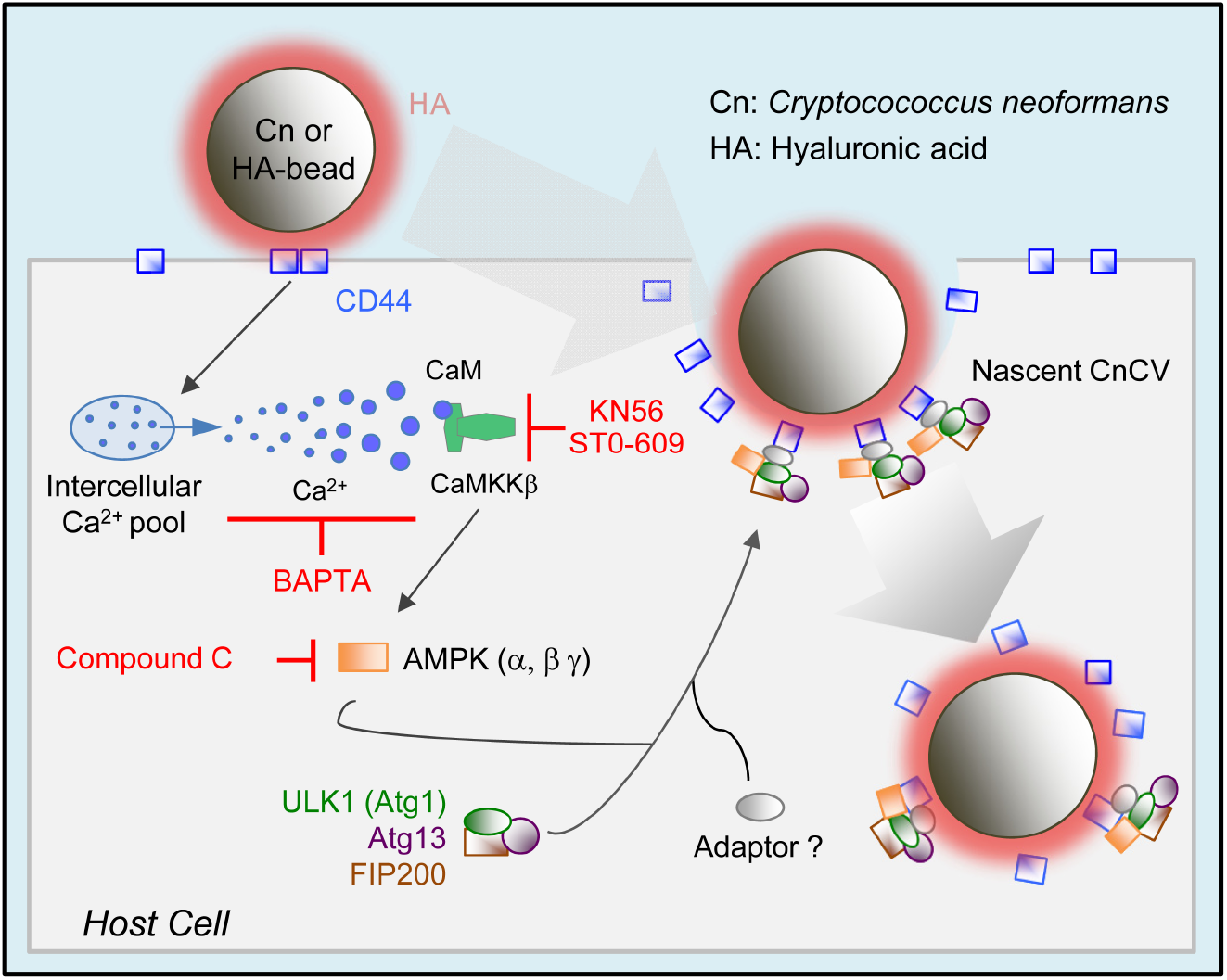

**In Brief:** Ding et al. reveal that interactions between fungal hyaluronic acid (HA) and host CD44 activate a Ca^2+^ - CaMKKβ-AMPK-ULK1 signaling pathway that recruits autophagy initiation complex components to forming phagosomes to drive fungal internalization.

**Highlights:**

- Fungal HA interactions with host cells drive a novel non-canonical, ligand-induced, autophagy pathway in phagocytic cells
- *Cryptococcus neoformans* recruits host CD44, together with AIC components and regulatory proteins, to forming phagocytic cups to initiate host cell internalization
- Fungal HA interactions with CD44 on host cell surfaces elevate intracellular Ca^2+^ concentrations, leading to activation of CaMKKβ
- A Ca^2+^-CaMKKβ-AMPK-ULK1 signaling axis is involved in HA and CD44 induced autophagy protein recruitment during Cn internalization

## Introduction

Autophagy is an orderly “self-eating” process in cells that coordinates the degradation of cellular components. Various types of autophagy have been described, including macrophagy, microphagy, mitophagy, chaperone-mediated autophagy and xenophagy (Galluzzi et al., 2017; Khandia et al., 2019; Kirkin and Rogov, 2019). Some pathogens subvert autophagic machinery to promote their intracellular survival and replication (Case et al., 2016; de Figueiredo and Dickman, 2016; de Figueiredo et al., 2015; Dickman et al., 2017; Pandey et al., 2018). Several signaling pathways control the onset, duration and outcome of autophagy induction in mammalian cells (Abada and Elazar, 2014; Rubinsztein et al., 2012). Pathways that include components of the autophagy initiation complex (AIC), including AMPK, ULK1, ATG13, FIP200 and ATG9, play important roles in these processes (Ganley et al., 2009; Hosokawa et al., 2009; Jung et al., 2009). For example, AMPK or ULK1 signaling can regulate ATG9 recruitment to nascent phagosome or autophagosome membranes (Mack et al., 2012). This recruitment process is believed to contribute to the elongation of the autophagosomal membrane (Mack et al., 2012).

*Cryptococcus neformans* (Cn) is a pathogen of global consequence that causes fatal fungal meningoencephalitis worldwide (Kozubowski and Heitman, 2012; Olszewski et al., 2010; Sabiiti and May, 2012). Cn is particularly pernicious in immunocompromised individuals, where lethal infection constitutes a significant risk (Warkentien and Crum-Cianflone, 2010). Cn can survive, replicate and persist in both intracellular and extracellular environments within mammalian hosts (Garcia-Rodas and Zaragoza, 2012). However, the molecular mechanisms that control intracellular parasitism remain poorly understood (Evans et al., 2018; Zaragoza, 2019). Towards addressing this issue, we reported a functional analysis of host factors that regulate the infection, intracellular replication, and non-lytic release of Cn from host cells (Qin et al., 2011). We extended these findings by performing a phosphoproteomic analysis of the host response to Cn infection (Pandey et al., 2017). This analysis demonstrated that host AIC proteins, and upstream regulatory molecules, contribute to the internalization and intracellular replication of the pathogen in macrophages (Pandey et al., 2017). This work also raised questions about the cellular and molecular mechanisms by which these proteins contribute to this phenotype.

The internalization of Cn into host cells is regulated, in part, by interactions between fungal components and host associated CD44, a major receptor for hyaluronic acid (HA) in mammalian cells (Jong et al., 2012; Jong et al., 2008). Moreover, CD44 has been shown to control phagocytosis of the pathogen (Jong et al., 2012; Jong et al., 2008). Interestingly, mice deficient for CD44 display reduced susceptibility to infection (Jong et al., 2012). The deficiency in phagocytosis accounts for this phenotype, leaving open the question of the mechanism by which CD44 controls fungal internalization. Here, we show that Cn phagocytosis by macrophages occurs by a novel mechanism whereby AIC proteins, including ULK1, ATG9 and ATG13, as well as the key upstream signaling component AMPKα, are recruited to forming phagosomes to promote the phagocytosis of the pathogen in a CD44-dependent fashion. Interaction of fungal HA and host CD44 activates a Ca^2+^-CaMKKβ (calcium/calmodulin-dependent protein kinase kinase β subunit)- AMPK-ULK1 signaling axis that supports Cn internalization into host cells. Taken together, our findings uncover unexpected roles for HA-CD44 interactions in conferring susceptibility to fungal infection and open up new avenues for therapeutic intervention for a fungal pathogen of global importance.

## Results

### Cn recruits host AIC components to forming Cn-containing phagosomes

To test the hypothesis that Cn infection of macrophages promotes the formation of a physical complex that contains AIC components, we used Forster Resonance Energy Transfer (FRET) imaging microscopy, which detects close molecular associations (< 10 nm) (Irving et al., 2014), to measure such interactions. The quenching of fluorescence in the donor fluorophore of a FRET pair accompanies the establishment of a close physical association between the pairs (Irving et al., 2014). We measured photon transfer between antibody labeled ATG13 and ULK1 or AMPKα and FIP200 in infected and uninfected RAW264.7 macrophages. We observed significant increases in the amount of FRET between these proteins in infected cells (**Figure S1A-D**). However, comparable FRET interactions were not observed in controls that were stained with a single label, or in uninfected samples (**Figure S1A-D**), thereby indicating the establishment of a close association between these proteins during infection.

### Recruitment of AIC components to forming phagosomes containing Cn is galectin 8 independent

Previous studies have shown that galectin 8, a β-galactoside-binding lectin, monitors endosomal and lysosomal integrity by binding to host glycans on damaged pathogen-containing vacuoles. This binding drives the ubiquitin-dependent recruitment of autophagy adaptor proteins (e.g., NDP52) and microtubule associated light chain kinase 3 (LC3), which in turn, promotes the recruitment of autophagosome biogenesis proteins to damaged pathogen-containing vacuoles (Thurston et al., 2012). Galectin 8 mediated autophagosomal targeting is relevant to the observed recruitment of AIC components to phagocytic cups because Cn containing vacuoles (CnCVs) are permeabilized after phagocytosis by macrophages (Johnston and May, 2010). Moreover, nascent CnCVs in infected macrophages recruit host LC3 (**Fig. S1E**) (Nicola et al., 2012; Qin et al., 2011), thereby implicating trafficking pathways that recruit LC3 to phagosomal membranes in controlling the intracellular lifestyle of Cn. With these ideas in mind, we tested the hypothesis that AIC recruitment to nascent CnCVs is associated with galectin 8 recruitment to these subcellular structures. We infected RAW264.7 macrophages that express a GFP-tagged variant of galectin 8 (GFP-Gal8) with Cn, and then used immunofluorescence microscopy (IFM) to determine whether GFP-Gal8 colocalized with AMPKα on nascent phagosomes containing Cn cells. We found that GFP-Gal8 did not display quantitative colocalization with either AMPKα or nascent phagosomes (**Figure S1F**). Consistent with these observations was the finding that UBEI-41, a potent and cell-permeable inhibitor of ubiquitin E1 activity (Yang et al., 2007), when added to macrophages at non-toxic doses (30 μM), did not diminish AIC recruitment to CnCVs (**Figure S1G**). Importantly, the inhibitor was washed out of the host cell culture media before Cn infection, thereby ensuring that the drug only targeted host cell components in these experiments. In addition, we found that nascent phagosomes decorated with the AIC component ULK1 colocalized with the endosomal marker EEA1 (**Figure S1H**). We also found that calreticulin, a Ca^2+^ binding ER (endoplasmic reticulum) protein, was recruited to phagocytic cups and nascent CnCVs (**Figure S1I**), and that LC3 displayed minimal colocalization with AIC components on nascent CnCVs (**Figure S1J, left**). CnCVs in B6J2 macrophages expressing dominant negative variants of the AIC regulatory component AMPKα recruited less LC3 (**Figure S1J, right**) and AIC components (**Figure S1K**), thereby suggesting that AIC recruitment to nascent Cn-containing phagosomes and LC3 recruitment to phagosome membranes could be morphologically and genetically dissected. Taken together, these findings indicated that recruitment of AIC to the nascent phagosomes and the induction of autophagy were galectin 8-independent events.

### A non-proteinaceous and/or non-capsular component controls AIC recruitment to nascent CnCVs

The observation that AIC and AIC regulatory components were recruited to forming phagosomes during Cn infection of macrophages encouraged us to determine the mechanism of recruitment. We infected host cells with live or dead Cn and found that fungal viability was not required for AIC recruitment because nascent CnCVs containing the heat-killed (HK) organism also efficiently recruited these proteins (**Figure 1A**). These observations were consistent with the hypothesis that non-proteinaceous, Cn-associated molecules activate the host AIC pathway.

**Figure 1.**
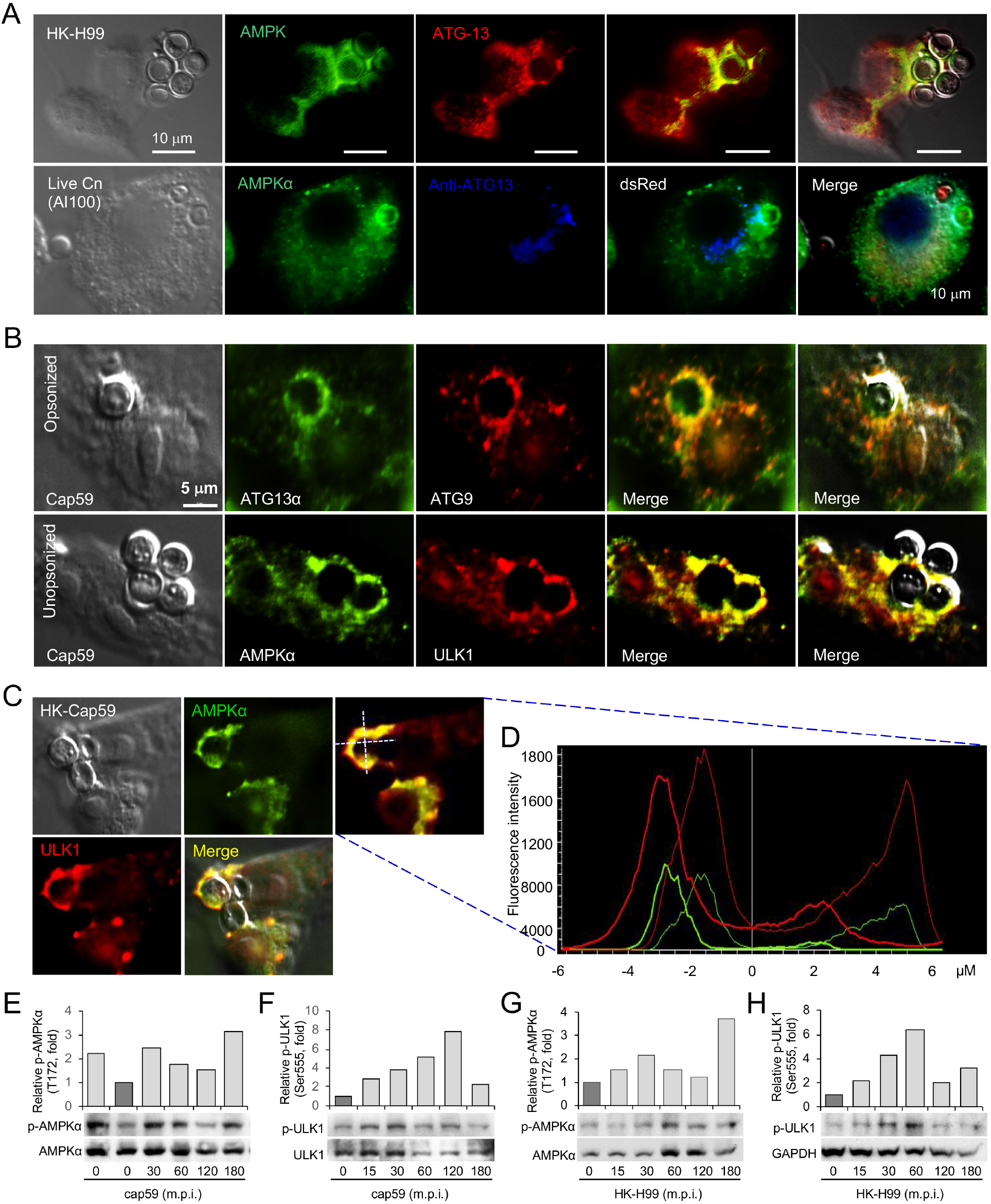
Non-proteineous components on Cn direct the recruitment of host AIC components to nascent phagosomes. (A, B) Colocalization of AMPKα and AIC components surrounding nascent CnCVs in host cells infected with live or heat-killed (HK) H99 (A) and opsonized or unopsonized acapsular cap59 strains (B) of Cn. (C, D) Recruitment of AMPKα and ULK1 to nascent CnCVs in host cells incubated with HK-cap59 at 3 h.p.i. (C) and the fluorescence intensity profile of AMPKα (green) and ULK1 (red) along the two crossed white lines (D). (E, F) Activation of host cell AMPKα (E) and ULK1 (F) by acapsular Cn strain cap59. (G, H) Activation of host cell AMPKα (G) and ULK1 (H) by HK Cn strain H99 (HK-H99).

Cn is encased in a carbohydrate-enriched capsule that is essential for intracellular parasitism and virulence (O’Meara and Alspaugh, 2012). To test the hypothesis that capsular components direct the recruitment of AIC and AIC regulatory proteins to forming phagosomes, we infected host cells with an acapsular mutant of Cn (Cap59), which displays defects in extracellular trafficking of glucuronoxylomannan (GXM) (Garcia-Rivera et al., 2004), and analyzed AIC recruitment to phagosomes containing cap59 strains. We found that forming and formed phagosomes that contained the acapsular strain also efficiently recruited AIC proteins (**Figure 1B-D, Figure S2A-C**). AIC components showed close associations with internalized Cn cells as detected by indirect immunofluorescence with anti-capsular monoclonal antibody 18B7 (Garcia-Rivera et al., 2004) (**Figure S2D**). Moreover, infection of host cells with live wild-type (WT) Cn resulted in the activation by phosphorylation of host cell ULK1 (Ser555), a key component of the AIC, and activation of the AIC regulatory protein AMPKα (Thr172) (Pandey et al., 2017). Similarly, the acapsular mutant and HK Cn also activated host AMPKα (Thr172) and/or ULK1 (**Figure 1E-H, Figure S2E**). However, when *S. cerevisiae* was incubated with host cells, similar AIC recruitment was not observed (**Figure S2F**), suggesting that phagocytosis of yeast cells by macrophages involves the participation of a distinct mechanism. Taken together, these findings suggested that Cn-specific, non-proteinaceous, non-capsular components activated the host AMPK-ULK1 signaling axis and promoted AIC recruitment to forming phagosomes.

### CD44-deficient host cells fail to recruit AIC proteins to forming phagosomes

CD44 regulates fungal internalization (Jong et al., 2012; Jong et al., 2008). This observation raised the intriguing possibility that CD44 interactions with other host cell components may also control the recruitment of AIC components to nascent CnCVs. To test this hypothesis, we first used fluorescence microscopy to determine whether CD44 colocalized with Cn cells or was recruited to forming or formed nascent CnCVs. We also tested whether CD44 deficient macrophages recruited AIC components to the nascent pathogen-containing phagosomes. We found that forming or formed phagosomes containing both capsular or acapsular strains displayed strong colocalization with CD44 (**Figure 2A**), and that CD44 was enriched at sites of contact between the pathogen and the host cell surface during a time course of infection (**Figure S2G**). Next, we tested whether CD44 colocalized with AIC regulatory AMPK and AIC components on nascent CnCVs. We found that AIC components, including ATG9 and ULK1, colocalized with CD44 on nascent CnCVs (**Figure 2B-C**). Compared to CD44^+/+^ bone-marrow derived macrophage (BMDM) controls, AMPKα showed reduced colocalization in CD44^−/−^ BMDMs (**Figure 2D**). As a result, Cn internalization in CD44-deficient cells was reduced (**Figure 2E-F**), indicating that host cell CD44 is required for Cn internalization, and that activation of the AMPK-ULK1 signaling axis and AIC recruitment is CD44-dependent.

**Figure 2.**
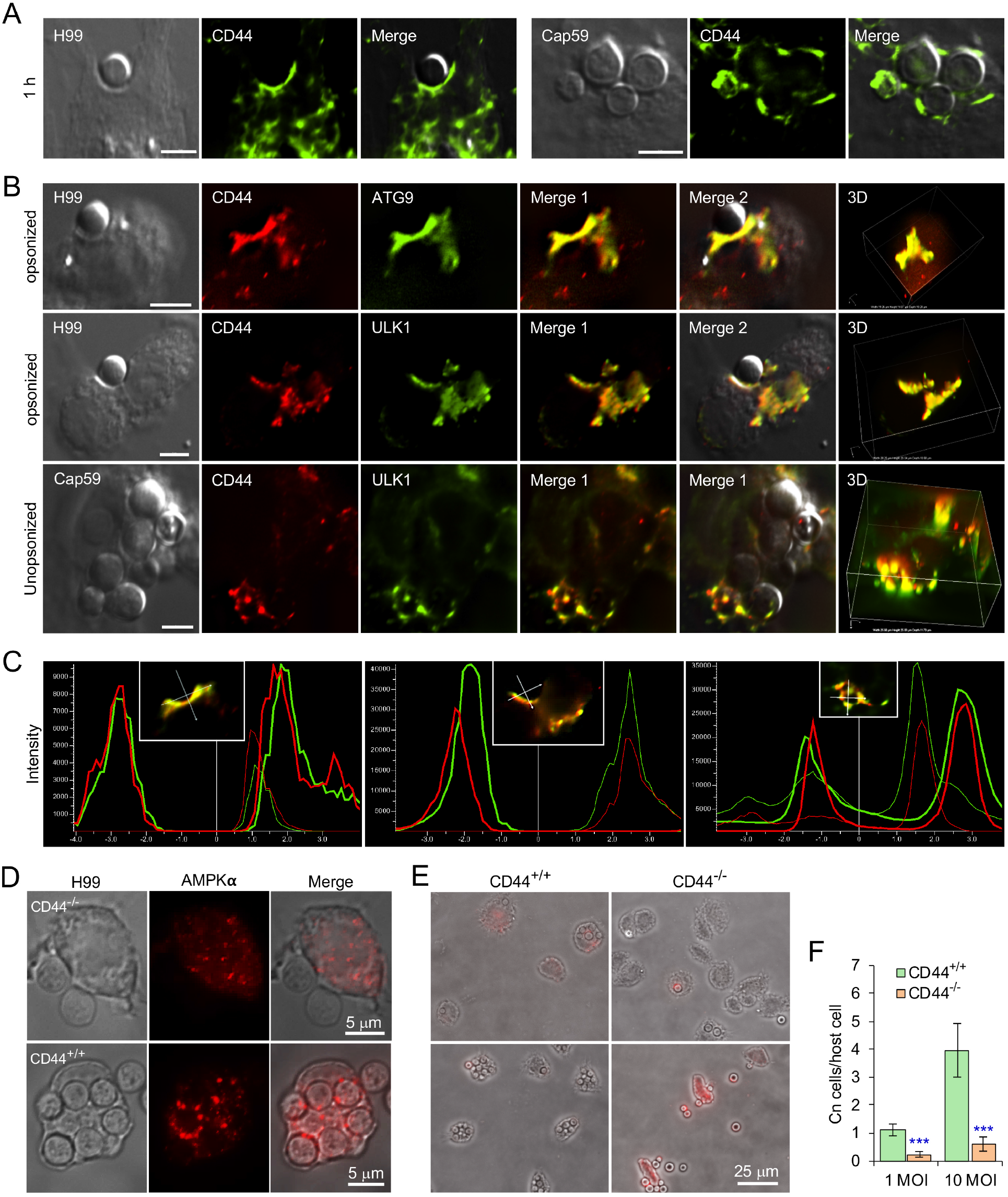
Host-associated CD44 is required for Cn host cell internalization. (A) Recruitment of host CD44 to nascent CnCVs that contained the wild-type (WT) strain H99 (leaf panel) or acapsular mutant strain cap59 (right panel) of Cn. At 1 hr post-infection (h.p.i.), host cells were fixed, permeabilized and processed for immunofluorescence microscopy using antibodies directed against the indicated host proteins. The antibody-stained samples were then subjected to confocal microscopy image analysis. (B) Colocalization of CD44 with the indicated AIC components in the vicinity of nascent CnCVs in host cells infected with the indicated Cn strains. (C) The fluorescence intensity profile of CD44 (red) and AIC components (ATG9 or ULK1) (green) along the two crossed white lines shown in the insets from (B, Merge 1). (D) AMPK recruitment to nascent CnCVs in CD44 knockout (KO, CD44^−/−^) and WT (CD44^+/+^) bone marrow-derived macrophages (BMDMs) at 3 h.p.i.. (E, F) Cn internalization in CD44 WT and KO BMDMs infected by Cn H99 assessed using image analysis approaches (E) and corresponding quantification (F) at 3 h.p.i.. Data represent the means ± standard deviation (SD) from three independent experiments. ***: significance at p < 0.001.

### *cps1* deletion mutants fail to recruit AIC components to nascent CnCVs

HA, a component of the Cn cell wall, is known to interact with CD44, an HA receptor on host cells, and to promote the internalization of the pathogen into human and murine brain microvascular endothelial cells (BMECs) (Jong et al., 2012; Jong et al., 2008). This observation raised the intriguing possibility that HA-CD44 interactions may promote the recruitment of AIC components to nascent CnCVs. To test this hypothesis, we first examined the internalization of Cn strains that harbor mutations in *cps1*, a gene that encodes hyaluronic synthase (Jong et al., 2007). Consistent with previous findings where Cn displayed reduced association with CD44-depleted murine BMECs compared to controls (Jong et al., 2012), we found that deletion of *cpsl* displayed reduced internalization of Cn into host macrophages (**Figure 3A-C, Figure S3A**). Next, we used fluorescence microscopy to determine whether *cps1*-deficient Cn strains recruited AIC components to nascent pathogen-containing phagosomes. We found the mutant strain displayed reduced recruitment compared to WT controls (**Figure 3D**). However, nascent CnCVs that contained strain C558 (*cps1*Δ::*CPS*1), in which the *cps1* mutation was complemented with a WT copy of the gene (Jong et al., 2008), displayed higher levels of AIC recruitment than their CPS1-deficient counterparts (**Figure 3D**). These data suggested that HA, the product of *cps1* activity, contributed to directing the recruitment of AIC components to nascent CnCVs, and implicated a role for CD44, the dominant HA receptor on macrophages, in regulating this process.

**Figure 3.**
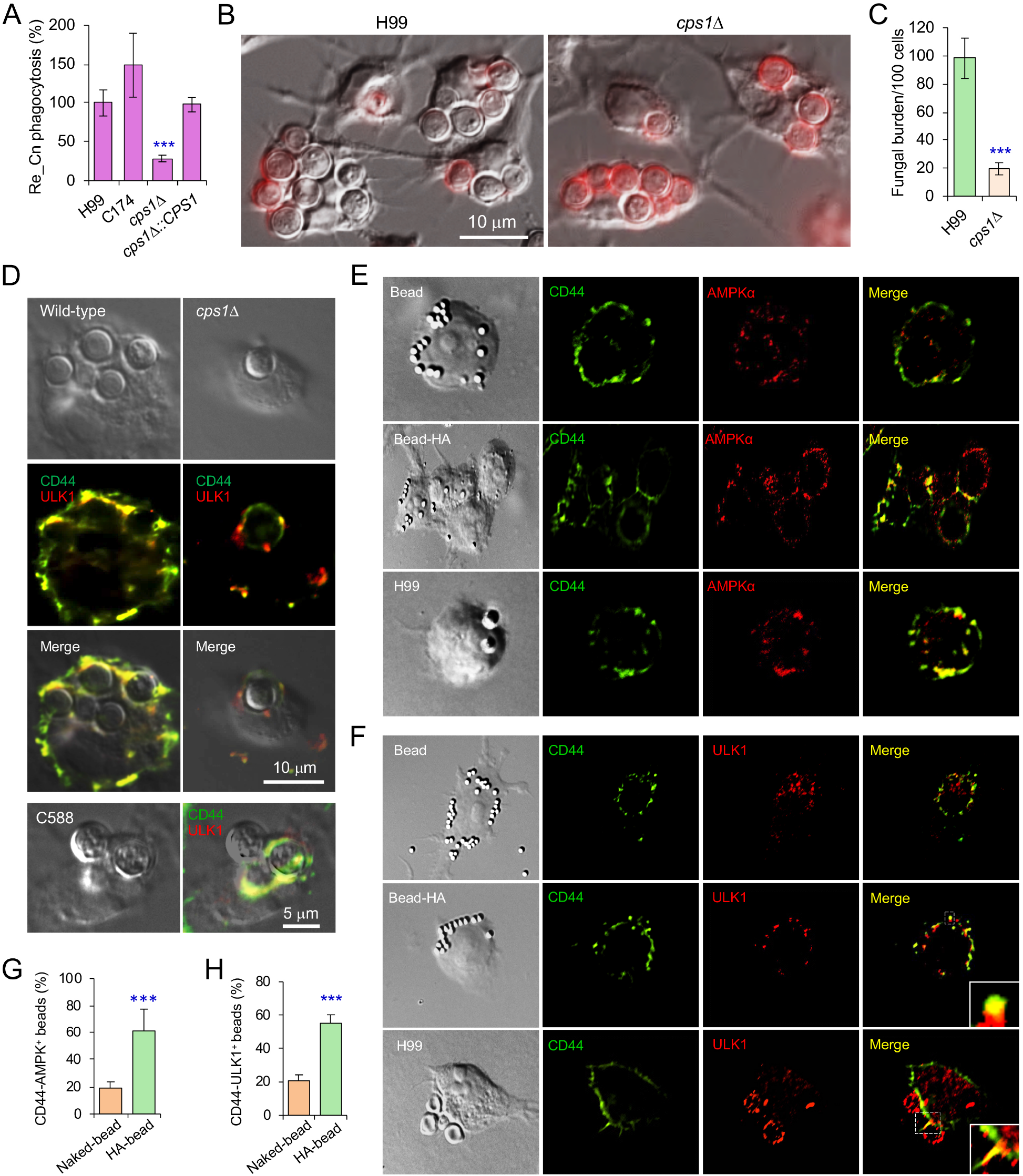
Fungal HA is required for recruitment of AMPK or AIC components to nascent phagosomes and Cn internalization. (A) Internalization of the wild-typ, *cps1*Δ and complemented Cn strains. (B-C) Internalization of the indicated Cn strains in BMDMs (B) and quantification of intracellular Cn cells (C) at 3 h.p.i.. (D) Confocal microscopy image analysis of recruitment of CD44 and the AIC component ULK1 by wild-type, *cps1*Δ, and complemented (C588) Cn strains at 3 h.p.i.. (E-F) Recruitment of host CD44 and AMPKα (E), or CD44 and AIC component ULK1 (F) to nascent phagosomes or CnCVs by HA-coated beads or Cn (H99), respectively. (G, H) Quantification of CD44 and AMPK positive beads (G) or CD44 and ULK1 positive beads based on confocal microscopy images as shown in (E-F). Data represent the means ± SD from three independent experiments. ***: significance at p < 0.001.

### Pathogen-derived HA drives interactions between CD44 and AIC components

To test whether HA was sufficient to induce CD44-mediated recruitment of AIC components to nascent CnCVs, we determined whether HA-coated beads induced similar recruitment to forming phagosomes. For these experiments, we covalently coupled HA to polystyrene beads and then incubated the HA-coupled beads with RAW264.7 macrophages for various lengths of time. We also used antibodies directed against AIC components in immunofluorescence microscopy experiments to visualize the recruitment of AIC components to forming phagosomes that contained beads. We found robust recruitment of AMPKα and AIC component ULK1 to forming phagosomes containing HA-coated beads (**Figure 3E-H**). We also incubated HA-coated beads with GFP-Gal8 expressing macrophages and found that recruitment of AMPKα or galectin 8 to the sites of bead internalization was not detected (**Figure S3B, upper**). However, co-localization of CD44 and AMPKα was observed to be colocalized with the HA-coated beads (**Figure S3B, lower**), Our observations therefore suggested that HA interactions with CD44 were necessary and sufficient to induce the formation of an AIC protein complex on forming phagosomes.

### Interaction of fungal HA with host CD44 activates AMPK and AIC pathways

The observation that interactions between HA and host CD44 recruited AIC components to forming phagosomes raised questions about the mechanism of AIC recruitment by these components. HA induces Ca^2+^ elevation in cells and may increase the activity of Ca^2+^-associated signaling pathways (Singleton and Bourguignon, 2002). For example, an increase of cytoplasmic Ca^2+^ levels can induce autophagy through CaMKKβ and AMPK pathways (Feng et al., 2020; Green et al., 2011). To test whether HA and CD44 interactions elevate intracellular Ca^2+^ ([Ca^2+^]_i_) levels and induce autophagy by Ca^2+^-mediated activation of CaMKKβ and AMPK pathways, we incubated host cells with Cn cells, HA, naked beads, or HA-coated beads, and then visualized [Ca^2+^]i levels and activation of CaMKKβ-AMPK pathways in the treated cells. We observed that compared to the controls, *cps1^+^-Cn* induced Ca^2+^-fluxes and increased [Ca^2+^]_i_ concentrations in a pulsed manner (**Figure 4A-C; Videos 1-2**). Similar results were observed in cells incubated with HA or HA-coated beads (**Figure S3C**), which is consistent with results from a previous report (Singleton and Bourguignon, 2002). Corresponding to the increase of [Ca^2+^]_i_ levels, activation of CaMKKβ and AMPKα was observed in H99 cells, (Pandey et al., 2017), or HA or HA-coated beads (**Figure 4D-E**). Importantly, phosphorylated AMPKα and ATG9 were coimmunoprecipitated from host cells incubated with HA-coated, but not naked, beads (**Figure 4F**).

**Figure 4.**
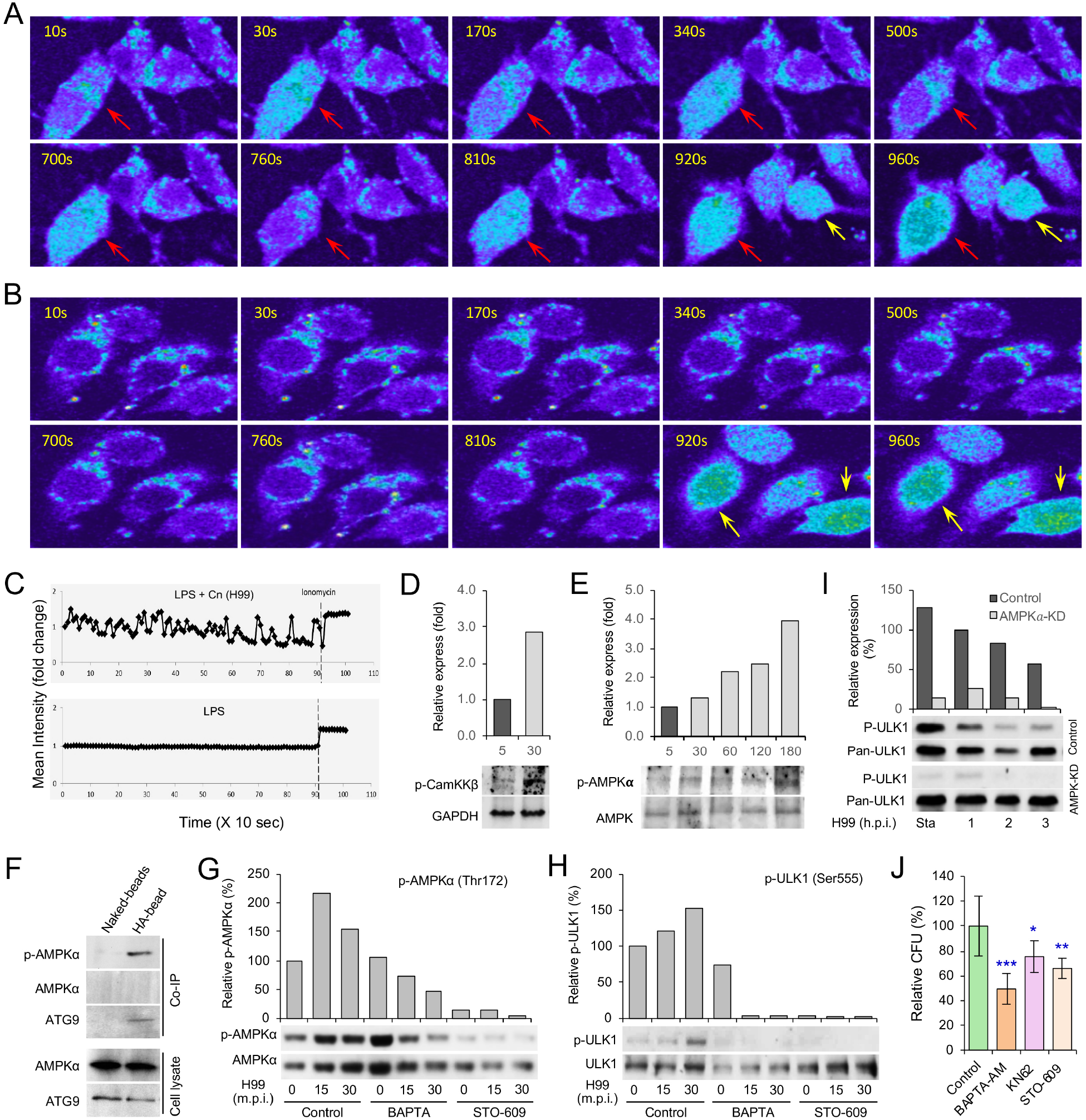
Cn infection activates CaMKKβ-AMPK-ULK1 signaling axis. (A, B) Cn infection results in intracellular Ca^2+^-flux in the infected host cells. Host cells incubated with *cps1^+^* Cn (A) or PBS (B) and the intracellular Ca^2+^ level was detected by excitation at 340 nm. Red arrows: Ca^2+^ releases from *cps1^+^*-Cn-infected host cells. Yellow arrows: Ca^2+^ releases from cells treated with ionomycin, a membrane permeable calcium ionophore used to increase intracellular calcium levels. (C) Mean Ca^2+^ flux intensity of Cn-infected (upper panel) and control (lower panel) cells. (D, E) Activation of host CaMKKβ (D) and AMPKα (E) in cells incubated with HA or HA-coated beads. (F) Co-immunoprecipitation (Co-IP) assays of host phosphorylated AMPKα and ATG9 trigged by HA-coated beads. Host cells were incubated with Cn. At 3 h.p.i., the infected cells were lysed and immunoprecipitated using antibodies against CD44. the precipitated material was then visualized by Western blot analysis using antibodies directed against p-AMPKα or ATG9. (G, H) Chelation of intracellular Ca^2+^ by BAPTA-AM and inhibition of the activity of CaMKKβ by STO-609 reduce activation of AMPKα (G) and ULK1 (H). (I) Depletion of AMPKα reduces activation of ULK1 during Cn internalization. Sta: starvation. (J) Cn internalization in host cells treated by the indicated drug via CFU assays. Data represent the means ± SD from three independent experiments. *, **, ***: significance at p < 0.05, 0.01, and 0.001, respectively.

To test whether activation of AMPK resulted from the increase of [Ca^2+^]_i_, we treated host cells with or without supplementation of assorted inhibitors, including BAPTA-AM (a [Ca^2+^]_i_-chelator), KN62 (a specific CaMK inhibitor), and STO-609 (a specific CaMKKα/β inhibitor), during infection with Cn. We found that host cells treated with these compounds displayed reduced activation of AMPK (**Figure 4G**); the activation of the downstream target ULK1 was also almost completely blocked (**Figure 4H**). During Cn infection, ULK1 is activated in a p-AMPK-dependent fashion (**Figure 4G-I**). Consequently, Cn internalization was reduced in cells in which [Ca^2+^]_i_ was chelated, the activities of CaMK or CaMKKα/β were inhibited or depleted (**Figure 4J; Figure S4A-D**), or AMPK or AIC components were depleted (**Figure S4E-F**). These findings suggest that interaction of HA with CD44 recruits AIC to forming phagosomes by release of Ca^2+^ and activation of the CaMKKβ-AMPK-ULK1 signaling axis.

## Discussion

The intracellular lifestyle of Cn is pivotal for pathogen colonization, dissemination and disease progression (Garcia-Rodas and Zaragoza, 2012; Johnston and May, 2013; Seider et al., 2010), as well as establishment of latent infection (Saha et al., 2007). Although the complete set of host factors that regulate intracellular parasitism remain obscure, published reports demonstrate that both “zipper” (receptor-mediated) and “trigger” (membrane ruffle dependent) mechanisms contribute to the internalization of Cn (Guerra et al., 2014). Moreover, interactions between the opsonized or unopsonized pathogen and host cell surface receptors are important to these processes (Shoham et al., 2001; Taborda and Casadevall, 2002). The findings reported here provide a new understanding of phagocytic mechanisms by demonstrating that fungal HA- and host CD44-dependent recruitment of AIC network components to nascent CnCVs play a central role in regulating the internalization of the fungus. Cn cells, opsonized or unopsonized and live or dead, as well as HA-coated beads, efficiently recruited host CD44, components of AIC and its regulatory protein AMPK to forming or formed phagosomes. As such, this report provides the first example of ligand- and receptor-induced recruitment of AIC proteins, including ULK1, FIP200, ATG13, and ATG9, to nascent phagosomes in macrophages.

Host cell galectin 8 is involved in defending against bacterial infection by recruiting the autophagic adaptor NDP52 to damaged *Salmonella-containing* vacuoles and in activating antibacterial autophagy (Thurston et al., 2012). Different from the defensive autophagy induced by galectin 8, significant recruitment of the danger receptor galectin 8 to forming or formed phagosomes during Cn internalization and/or intracellular replication was not observed, suggesting that a different strategy is employed to recruit elements of the autophagy machinery through interactions between fungal HA and host cell CD44. Besides AMPK and AIC components, the endosomal and lysosomal markers EEA1, M6PR, and Cathepsin D, as well as the ER marker calreticulin, were recruited to nascent phagosomes (Qin et al., 2011). Interestingly, co-localization of the endocytic marker EEA1 and the AIC component ULK1 was also observed, suggesting that the endosomal and lysosomal pathways, ER-derived membrane and selective autophagy machinery are involved in the internalization of Cn into host cells. How these factors coordinate this process and the involvement of other ubiquitin-binding autophagic adaptors related to the AIC remain to be characterized.

AMPK can be activated by cellular stresses that elevate AMP levels by means of allosteric binding of AMP to sites in the g subunit AMPK, and by phosphorylation of Thr172 in AMPKα by the tumor suppressor LKB1, CaMKKβ, or the transforming growth factor-β-activated kinase (TAK1) (Hardie et al., 2012; Kola et al., 2006; Zadra et al., 2015). Activation of AMPK can induce autophagy via direct phosphorylation of ULK1 (Egan et al., 2011; Kim et al., 2011; Zhao and Klionsky, 2011). Previous observations showed that HA interactions with CD44 induces Ca^2+^ elevation in cells (Singleton and Bourguignon, 2002), and that increases in cytoplasmic Ca^2+^ concentrations induce autophagy through activation of CaMKKβ and AMPK pathways (Feng et al., 2020; Green et al., 2011). Similarly, our findings demonstrate that infection by Cn recruits host cell CD44 to CnCVs, induces intracellular Ca^2+^-flux in the infected host cells, activates CaMKKβ and AMPK, and recruits AIC components to forming or nascent CnCVs. Cn infection also activates LKB1, and depletion of LKB1 reduced Cn internalization into host cells (Pandey et al., 2017). However, how the interaction of HA and CD44 activates AMPK through the upstream regulatory proteins LKB1 and TAK1 remains to be further characterized.

Our data support a stepwise model in which several sequential molecular events control the internalization of Cn in macrophages. First, interactions between fungal HA and CD44 on the surface of host cells stimulates the release of Ca^2+^ and elevates intercellular Ca^2+^ levels, which results in the activation of CaMKKβ and AMPK. Second, phosphorylation of ULK1 occurs in an AMPK-dependent fashion. Third, AMPK-dependent activation of ULK1 recruits AIC components, including the ULK1-ATG13-FIP200 complex, ATG9, and LC3, to forming phagocytic cups containing the fungus. Fourth, the coordinated activities of AIC components drive the internalization of the pathogen into host cells. Finally, AIC component interactions with CnCVs gradually diminish as the pathogen establishes a replicative niche in host cells (**Figure for graphical Abstract**).

Finally, it is notable that several proteins in AMPKα and AIC regulatory networks are targets of commonly prescribed drugs (e.g., AMPK, a target for metformin) or under development as targets for pharmaceutical intervention (e.g., ULK1) (Egan et al., 2015). Several fungi, including *Candida* spp. and *Histoplasma capsulatum,* are capable of intracellular parasitism (Garcia-Rodas et al., 2011; Howard, 1965; Woods, 2003). Therefore, our findings may open up new therapeutic possibilities for preventing cryptococcosis and other infections caused by intracellular pathogens.

## Acknowledgement

We gratefully thank Arturo Casadevall (Department of Microbiology and Immunology, Albert Einstein College of Medicine, Yeshiva University) for anti-cryptococcal antibodies, and Steve Fullwood and Kalli Landua (Nikon Instruments) for expert assistance with the microscopy analysis. This work is supported by the Texas AM Clinical Science Translational Research Institute Pilot Grant CSTR2016-1, DARPA (HR001118A0025-FoF-FP-006), NIH (R21AI139738-01A1, 1 R01AI141607-01A1, 1R21GM132705-01), the National Science Foundation (DBI 1532188, NSF0854684) and the Bill Melinda Gates Foundation, the Defense Advanced Research Projects Agency (Agreement HR001118A0025-FoF-FP-006) to PdF; NIH grant awards NIH 1R01 AI48496-01A1 and NIH 1U54AI057156-0100 to TAF; the National Natural Science Foundation of China (# 81371773) to QMQ. Any opinions, findings, and conclusions or recommendations expressed in this material are those of the author(s) and do not necessarily reflect the views of the funding agencies.

## Author Contributions

P.D., Q.M.Q., T.A.F. designed the experiments. S.L.D., A.P., X.H.F., Q.M.Q, J.Y., L.F.d.C., R.M. S.L.B. performed experiments. P.D., T.A.F., A.R., R.M., R.O.W., K.P. provided reagents/analysis tools. P.D., Q.M.Q., S.L.D., X.H.F., A.P., T.A.F. analyzed data. P.D., Q.M.Q. supervised the work and wrote the manuscript.

## Star Methods

### KEY RESOURCES TABLE

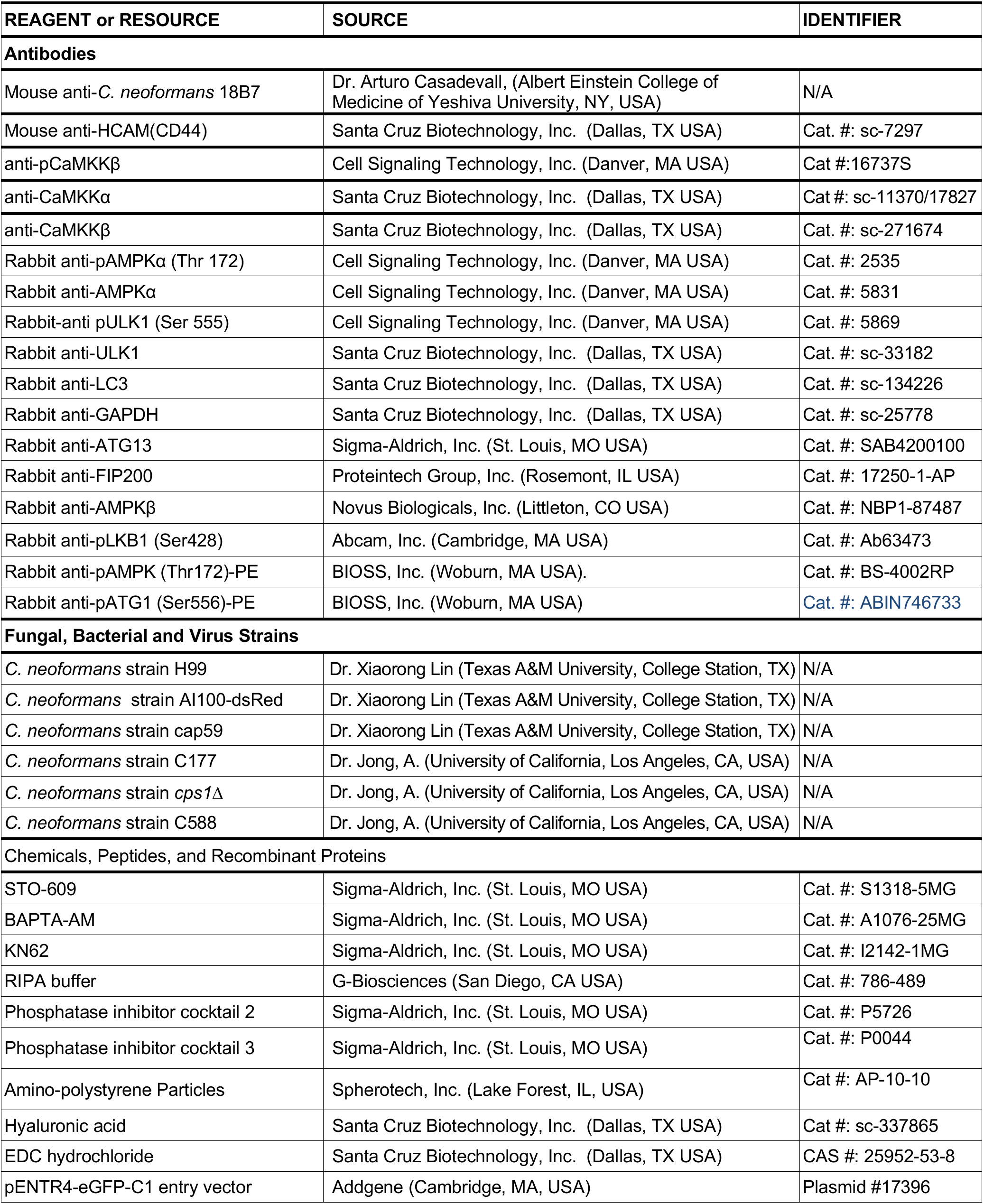
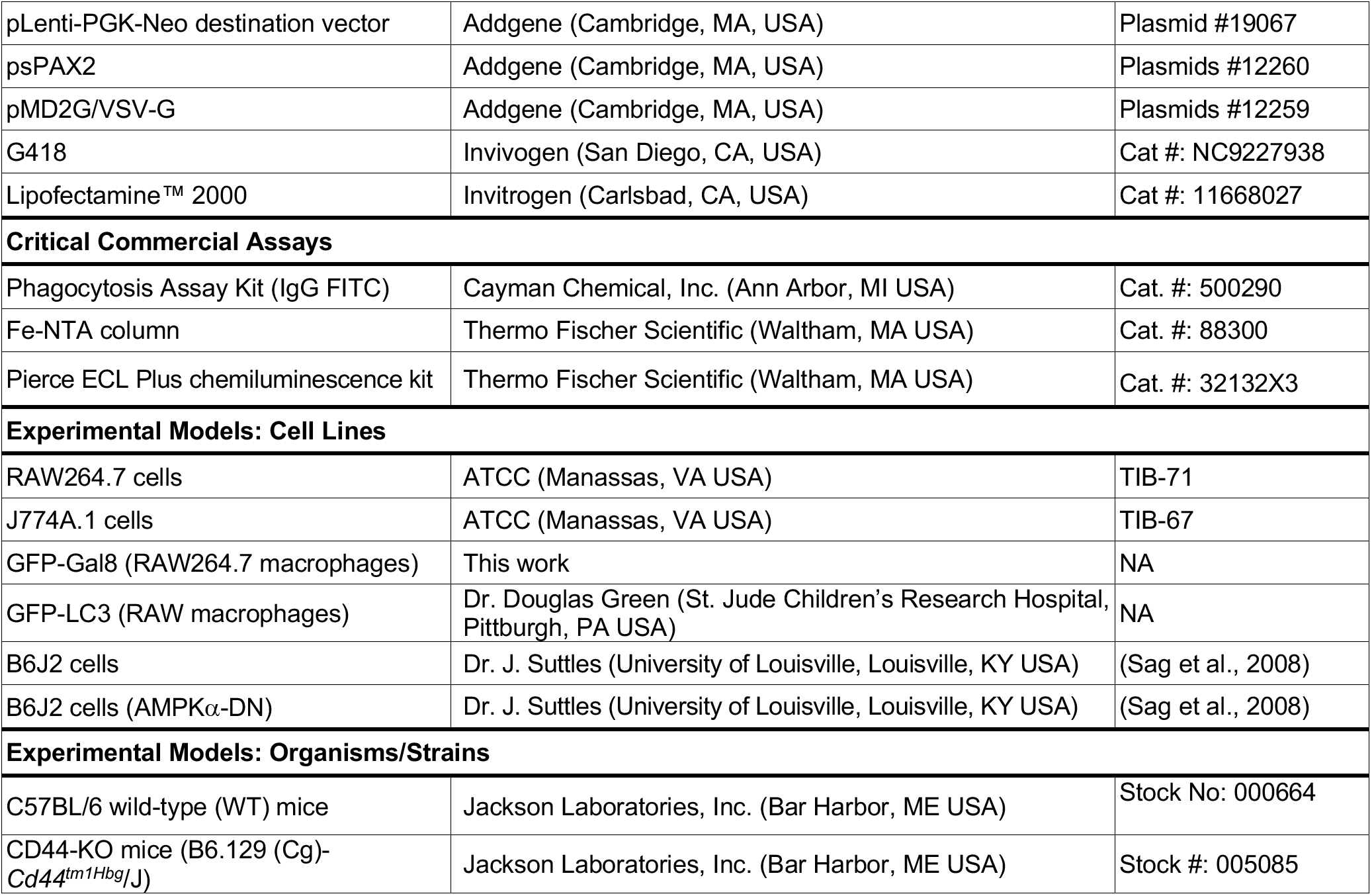

### CONTACT FOR REAGENT AND RESOURCE SHARING

Further information and requests for resources and reagents should be directed to and will be fulfilled by the Lead Contact, Paul de Figueiredo.

### EXPERIMENTAL MODEL DETAILS

#### Bone marrow-derived macrophage harvest and cultivation

Bone marrow cells were collected from the femurs of littermate control and CD44 KO mice, and cultivated in L929-cell conditioned media [DMEM medium containing 20% L929 cell supernatant, supplemented with 10% (v/v) FCS, penicillin (100 U/ml), and streptomycin (100 U/ml)]. After 3 days of culture, non-adherent precursors were washed away and the retained cells were propagated in fresh L929-cell conditioned media for another 4 days. For experimentation, BMDMs were split in 24-well plates at a density of 2.5 × 10^5^ cells per well in L929-cell conditioned media and cultured at 37°C with 5% CO_2_ overnight before use.

#### Method Details

##### *Cryptococcus* Strains, Cell Culture, *Cryptococcus* Infection, and CFU Assay

Yeast forms of Cn cells were cultured on YPD (Difco™) agar plates and maintained on the plates for 4 to 5 days prior to experimentation. Mammalian cell lines were routinely incubated in DMEM supplemented with 10% FBS in a 5% CO_2_ atmosphere at 37°C. Preparation of cryptococcal and host cells for infection as well as CFU assays that measured internalized Cn cells were performed as previously described (Qin et al., 2011; Pandey et al., 2017).

##### Immunofluorescence Microscopy Assays for Cn Phagocytosis

Mammalian host cells were first cultivated on 12 mm glass coverslips (Fisherbrand) on the bottom of 24-well plates (Falcon) for 12-16 h before infection. Next, the host cells were infected with the tested Cn cells and incubated at 37°C in a 5% CO_2_ atmosphere. At the indicated time points postinfection, culture media was removed and the infected host cells were washed 6 to 8 times with PBS (pH 7.4) before being fixed with 3.7% formaldehyde. The fixed cells were then processed Cn phagocytosis assays using antibody (18B7) directed against Cn as previous described (Qin et al., 2011; Pandey et al., 2017) without adding Triton X-100 to the staining buffer. The numbers of internalized (unstained) and extracellular (stained) Cn cells were then quantified and plotted. Images represent a representative image from triplicate replicates, with 100 fields imaged per replicate.

##### Drug/compound Treatments

Murine macrophage J774.A1 or RAW264.7 cells were overnight cultured in 48 well plates and then coincubated with assorted pharmacological compounds, including KN62, STO-609 or BAPTA-AM for 3 h. The media containing the compounds was then removed. For CFU assays, the drug-treated cells were extensively washed with fresh medium and then infected with Cn cells. At 3 h.p.i., the infected cells were lysed and performed CFU assay as previously described (Qin et al., 2011; Pandey et al., 2017). For western blots, the compound-treated cells were washed 3 times with cold 1 × PBS, pH7.4, lysed and performed immunoblotting assay as previously describe (Pandey et al., 2017).

##### Lentivirus-Mediated Depletion of Host Proteins

The pSuperRetro retrovirial vector system (OligoEngine, Inc.) was used to knockdown target gene expression in murine cells according to the manufacturer’s instructions. The oligonucleotides used for shRNA construction to knockdown the expression of mouse genes and the accompanying references are listed in **Table S1**. Transfection was performed in 6-well plates containing 1.5 × 10^5^ RAW264.7 or B6J2 cells. Clones with the insert stably integrated were selected with puromycin. Western blot was performed to validate the depletion of the targeted proteins. All Westerns were performed in triplicate and representative findings are shown.

##### Generation of GFP-tagged Gal8 RAW264.7 cells

The GFP-galectin-8 expression construct was made by first cloning the cDNA sequence of *Lgals8* (from RAW 264.7 cells) into the pENTR4-eGFP-C1 entry vector (Campeau et al., 2009), resulting in a fusion of eGFP on the N-terminus of galectin-8. This construct was fully Sanger sequenced (Eton Biosceinces, San Diego, CA) to verify the fusion protein was in-frame and error-free. GFP-Gal8 was then Gateway cloned with LR Clonase (Invitrogen) into the pLenti-PGK-Neo destination vector (Campeau et al., 2009). Lenti-X 293T cells (Takara Bio) were co-transfected with pLenti-GFP-Gal8 and the packaging plasmids psPAX2 and pMD2G/VSV-G (Addgene Plasmids #12259-60) to produce lentiviral particles. RAW 264.7 cells were transduced with GFP-Gal8 lentivirus for two consecutive days plus 1:1000 Lipofectamine 2000 (Invitrogen) and selected for 5 days with 750 μg/ml G418 (Invivogen). Expression of GFP-Gal8 was confirmed by Western blot analysis and fluorescence microscopy.

##### Confocal Microscopy Assays

Host cells were infected with live or heat killed, opsonized or unopsonized capsular or acapsular strains of Cn. At the indicated times post-infection, host cells were fixed for confocal immunofluorescence microscopy analysis using antibodies directed against the indicated host proteins. Immunofluorescence microscopy staining and imaging methods (Qin et al., 2008; Qin et al., 2011; Pandey 2017; Pandey 2018) were used to determine the subcellular localization of host AIC components in infected host cells. Samples were observed on a laser scanning confocal microscope or on a confocal fluorescence microscope (ECLIPSE Ti, Nikon). Confocal images (1,024 × 1,024 pixels) were acquired and processed with NIS elements AR 3.0 software (Nikon) and assembled with Adobe Photoshop CC 2019 (Adobe Systems, CA, USA). Digital image analysis and quantification was performed as previously described (Qin et al., 2011). Findings from our subcellular localization analyses were not an artifact of secondary antibody cross reactivity with host or pathogen components because negligible fluorescence signal was observed when infected cells were stained with secondary antibodies alone, or the pathogen alone was stained with the antibodies used in the experiments (**Figure S4G**).

##### Protein pull-down assay with HA-coated or naked beads

Spherical amino polysterene beads (size: 1.0-1.4 μm, Spherotech Inc, IL, USA) were covalently coupled using EDC [*N*-Ethyl-*N*’-(3-dimethylaminopropyl)carbodiimide hydrochloride] to hyaluronic acid (HA) (Santa Cruz Biotec.) as per the manufacturer’s (Spherotech) instruction. Briefly, 200 μl of 0.05 M sodium acetate buffer (pH 5.0), 2 mg of HA, 2 ml of 5% w/v Amino particles and 20 mg of EDC were mixed in a glass centrifuge tube. The contents were vortexed and incubated for 2 hrs at ambient temperature on a rotary mixer. Following incubation, the tube was centrifuged at 3000 × g for 15 minutes and supernatant was carefully discarded. The pellet was washed twice in 1 × PBS and resuspended in 2 ml of 1 × PBS to obtain 5% w/v suspension of HA coated beads. RAW 264.7 cells (3×10^5^) were washed with 1 × PBS. Next, a bead incubation solution containing 5 μl of HA-coated or naked beads in 200 μl of PBS was added to the washed cells. The cells were centrifuged for 10 min at 1500 RPM and then incubated in a humidified incubator containing 5% CO_2_ for 2 hrs. The treated cells were washed 2-4 times with ice-cold 1 × PBS and lysed in 80 μl RIPA buffer supplemented with a cocktail of protease and phosphatase inhibitors. Beads were separated from the lysate by high-speed centrifugation at 4°oC. The separated beads were washed 3 times with RIPA buffer and finally resuspended in 80 μl RIPA buffer and 20 μl of 5 × sample buffer (Thermo Scientific) and boiled for 5 mins. Samples were finally resolved by SDS-PAGE and subjected to western blot analysis.

##### Immunoblotting Analysis

Preparation of protein samples and western blot analysis were performed as described previously (Qin et al., 2011; Pandey et al., 2017; Pandey et al., 2018). Blot densitometry was performed using the ImageJ (http://rsbweb.nih.gov/ij/) software package. All Westerns were performed in triplicate and representative findings are shown.

##### Quantification and Statistical Analysis

The quantitative data presented in this work represent the mean ± standard deviation (SD) from at least three independent experiments. To easily compare results from independent experiments, the data from controls, such as protein expression level, blot densitometry, CFU, intracellular Cn number, etc., were normalized as 1 or 100%. The significance of the data was assessed using the Student’s *t*-test to assess statistical significance between two experimental groups or a one-way ANOVA test to evaluate the statistical differences of multiple comparisons of the data sets.

## Supplemental Figures and Figure Legends

**Figure S1.**
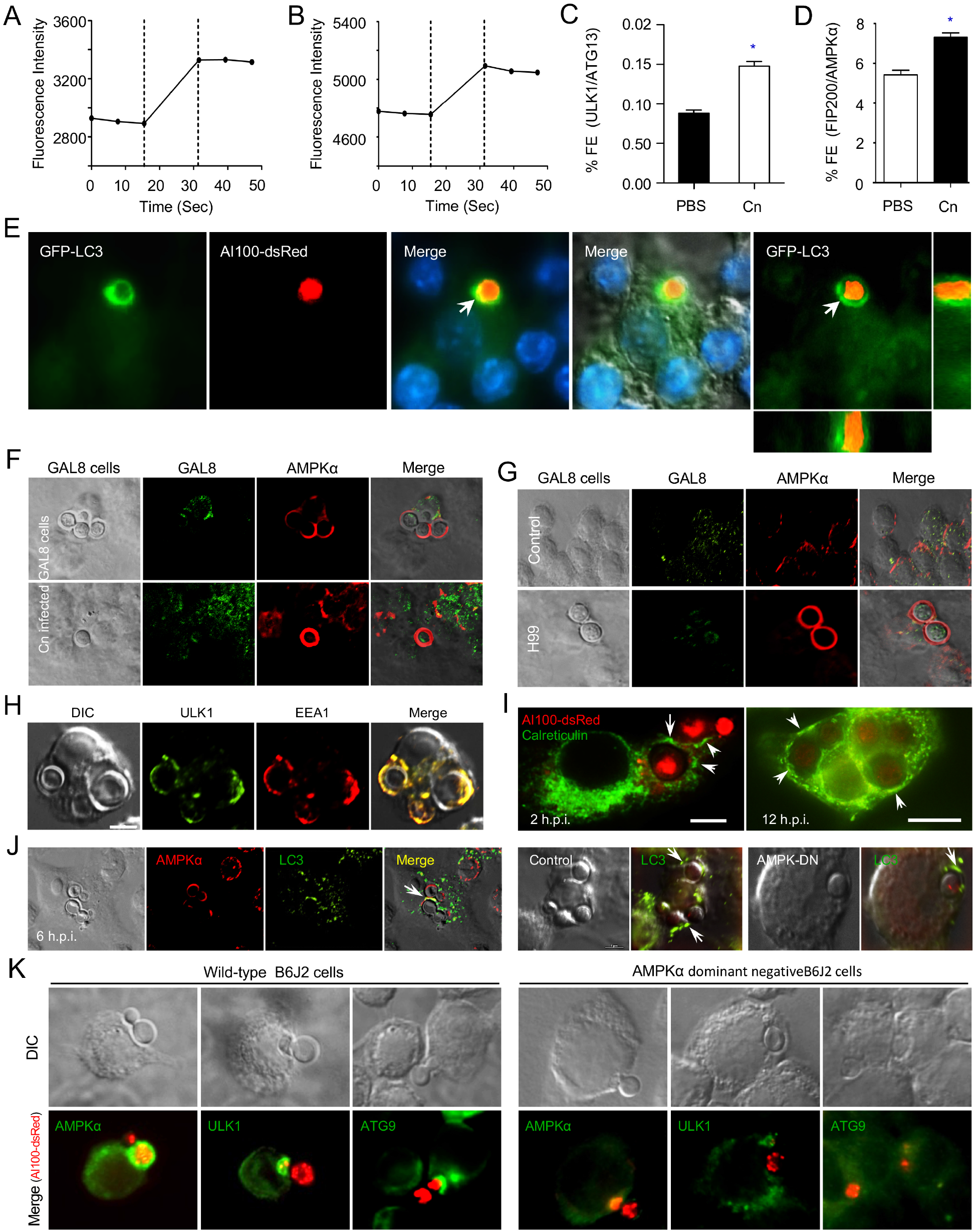
Recruitment of AIC components, but not galection 8, to nascent phagosomes during Cn internalization. (A, B) Fluorescence intensity of ULK1 and ATG13 (A) or FIP200 and AMPKα (B) during Cn internalization. (C, D) Interactions between host ULK1 and ATG13 (C) or FIP200 and AMPKα (D) recruited to Cn-containing vacuoles (CnCVs) in RAW264.7 macrophages. The infected host cells were quantified using FRET analysis. Data represent the means ± standard deviations (SD) from at least three independent experiments. * indicates significance at P < 0.05. (E) Recruitment of host LC3 to the nascent CnCV s during Cn internalization. (F, G) Host galectin 8 is not recruited to nascent phagosomes containing Cn cells during Cn internalization by host cells expressing GFP-galectin 8 (F) or cells supplemented with UBEI-41, a cell permeable inhibitor of ubiquitin-activating enzyme E1 (G). (H) Colocalization of host early endosomal marker EEA1 with AIC component ULK1 surrounding nascent CnCVs. (I) Recruitment of host endoplasmic reticulum marker calreticulin to nascent and formed CnCVs. (J) Recruitment of host AMPKα and LC3 to nascent phagosomes containing Cn cells in infected bone marrow derived macrophages (BMDMs, left panel) or in B6J2 macrophages expressing dominant negative variant of AMPKα (AMPK-DN) and control (right panel). (K) Less recruitments of host AMPKα and AIC component ULK1 or ATG9 to nascent CnCVs in B6J2 macrophages expressing dominant negative AMPKα variant during Cn internalization.

**Figure S2.**
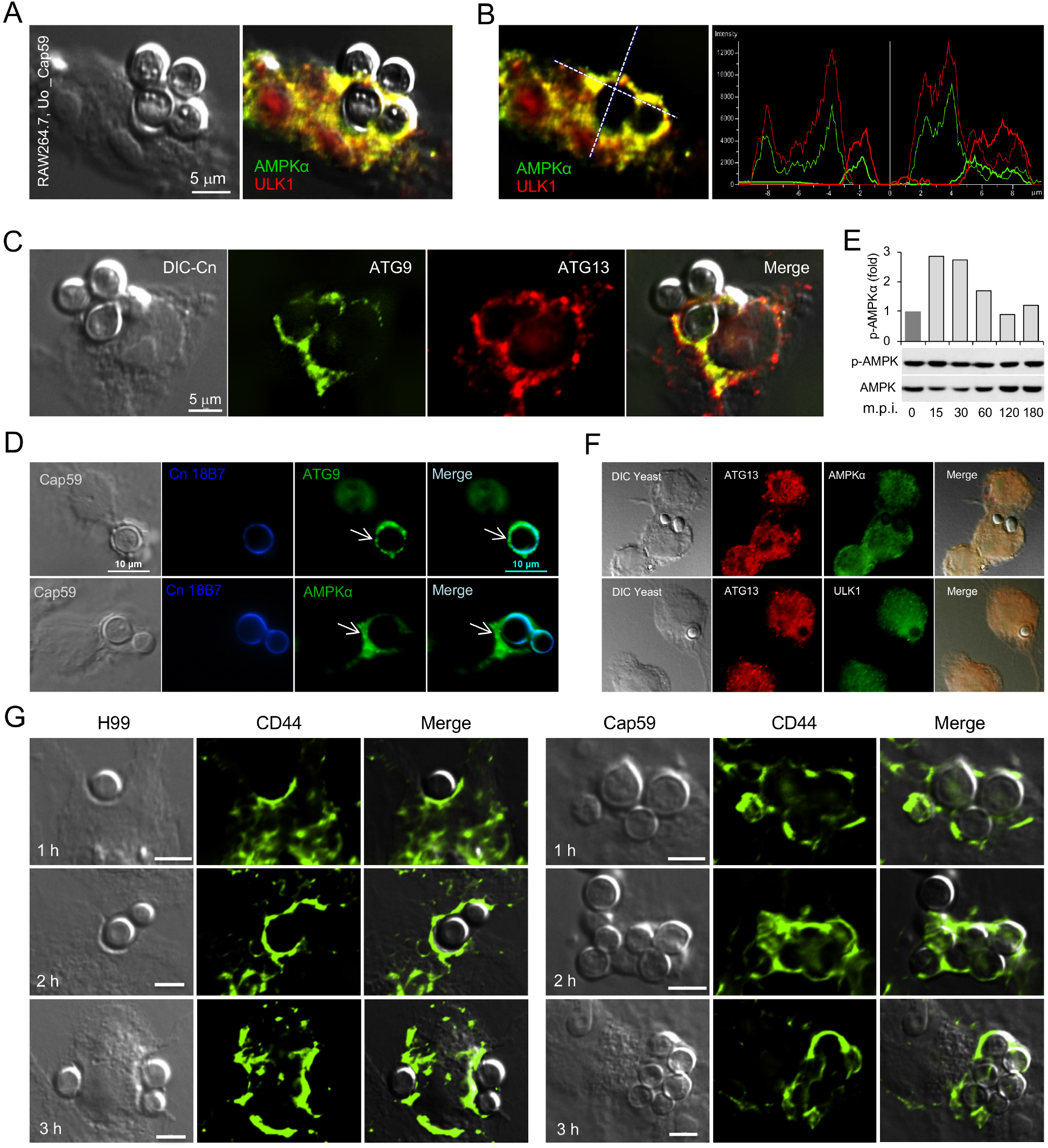
Recruitment of host AIC components and CD44 to forming or nascent CnCVs during Cn internalization. (A) Recruitment of AMPKα and AIC components ULK1 to nascent CnCVs. (B) The fluorescence intensity profile of AMPKα (green) and ULK1 (red) along the two crossed white lines (left panel). (C) Colocalization of ATG9 with ATG13 in the vicinities of CnCvs. (D) Colocalization of host AMPKα or ATG9 with cryptococcal glucuronoxylomannan (GXM)-specific monoclonal antibody 18B7. (E) Activation of hos AMPKα by heat-killed (HK) Cn cells (cap 59) during phagocytosis. Representative results from one of three independent experiments are shown. (F) Recruitment of AIC components and AMPKα to nascent phagosomes is hardly detected during phagocytosis of yeast *(Saccharomyces cerevisiae)* cells by host cells. (G) Recruitment of CD44 during a time course (3 hr) of Cn internalization.

**Figure S3.**
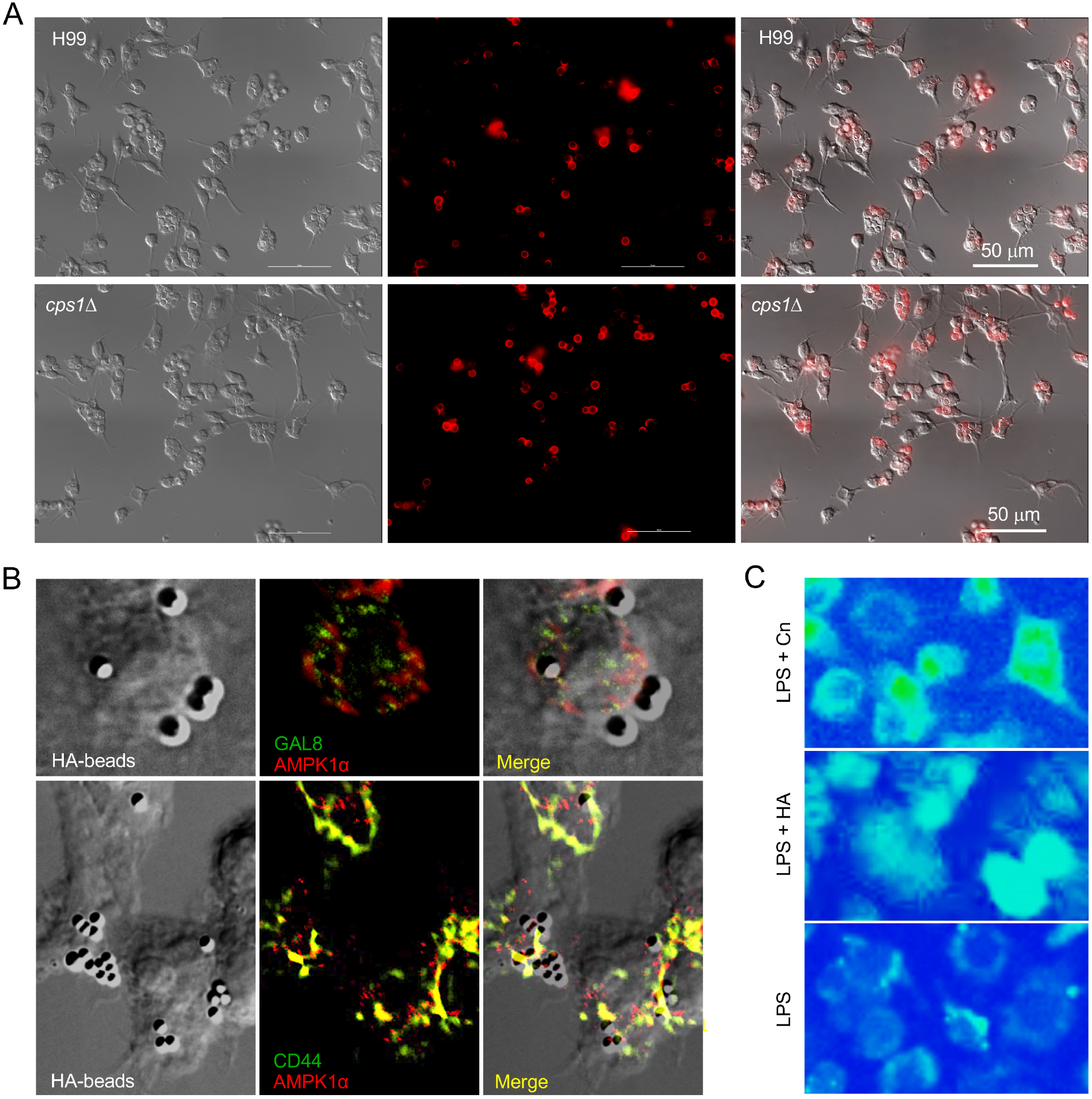
Fungal hyaluronic acid (HA) is required for Cn internalization. (A) Disruption of Cn *cpsl* impairs internalization of the fungal pathogen (related to Figure 3B-C). (B) Recruitment of CD44, not galectin 8, to the forming or nascent phagosomes associated with HA-coated beads. Upper panel: localization of galectin 8 and AMPKα in cells incubated with HA-coated beads. Low panel: colocalization of CD44 and AMPKα to the forming or nascent phagosomes related to HA-coated beads. (C) Incubation of host cell with Cn harboring *cpsl* (top) or HA (middle) increases intracellular Ca^2+^ level (exciated at 340 nm).

**Figure S4.**
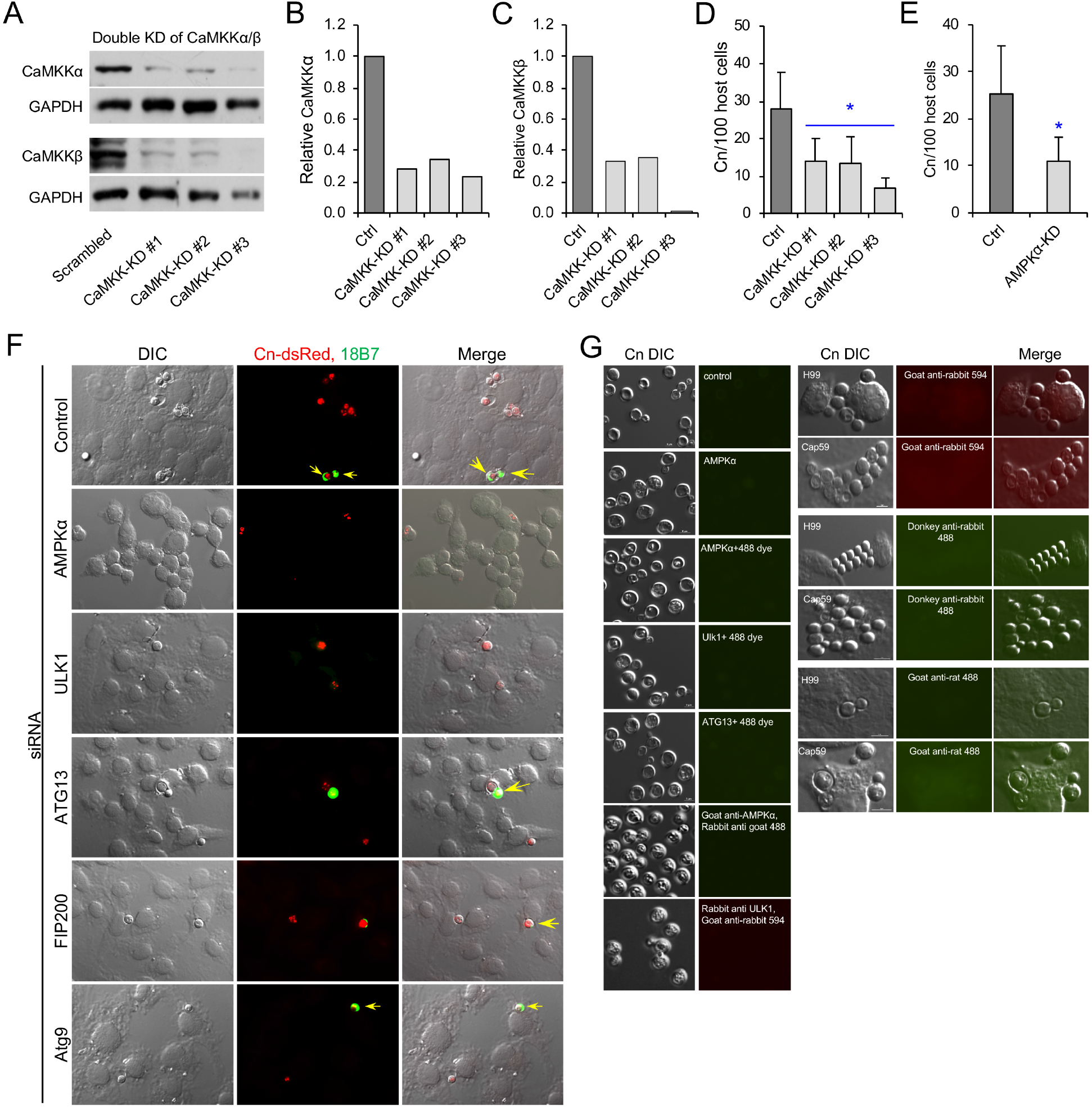
Components in the CaMKK -AMPK-ULK1 signal aix play important roles in Cn internalization. (A) Immunobloting assay to test the depletion of CaMKKα/β protein level by the shRNA approach. (B, C) Relative protein expression levels of CaMKKα (B) or CaMKKβ (C) in RAW macrophages transfected with scambled or CaMKKα/β shRNAs. Data from a representative experiment shown in (A). (D, E) Depletion of host cell CaMKKα/β (D) or AMPKα (E) reduces Cn internalization determined by immunofluorescence microscopy assay. Data represent the means ± SD from three independent experiments. *: significance at p < 0.05. (F) Depletion of host cell AMPKα and AIC components reduces Cn internalization determined by immunofluorescence microscopy assay. Images from a representative experiment of three independent experiments. (G) Cn cells harvested from infected host cells (leaf panel) or during host infection (right panel) do no display cross activities with the primary or secondary antibodies used in this work.

**Table S1.**
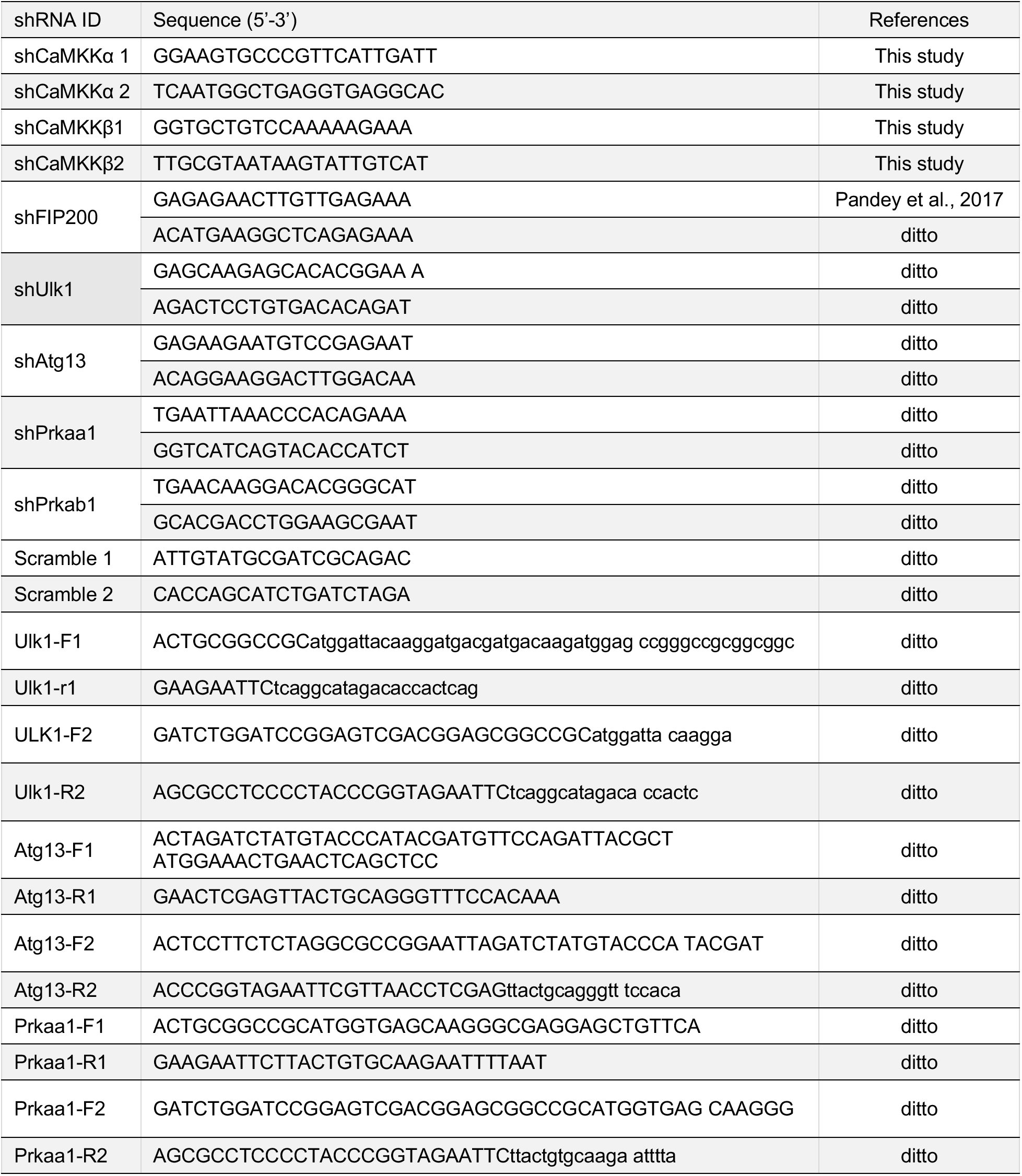
shRNAs and PCR primers used in this study

