## Supplemental Videos 1 and 2 for "Interactions between fungal hyaluronic acid and host CD44 promote internalization by recruiting host autophagy proteins to forming phagosomes"

### Slide 1
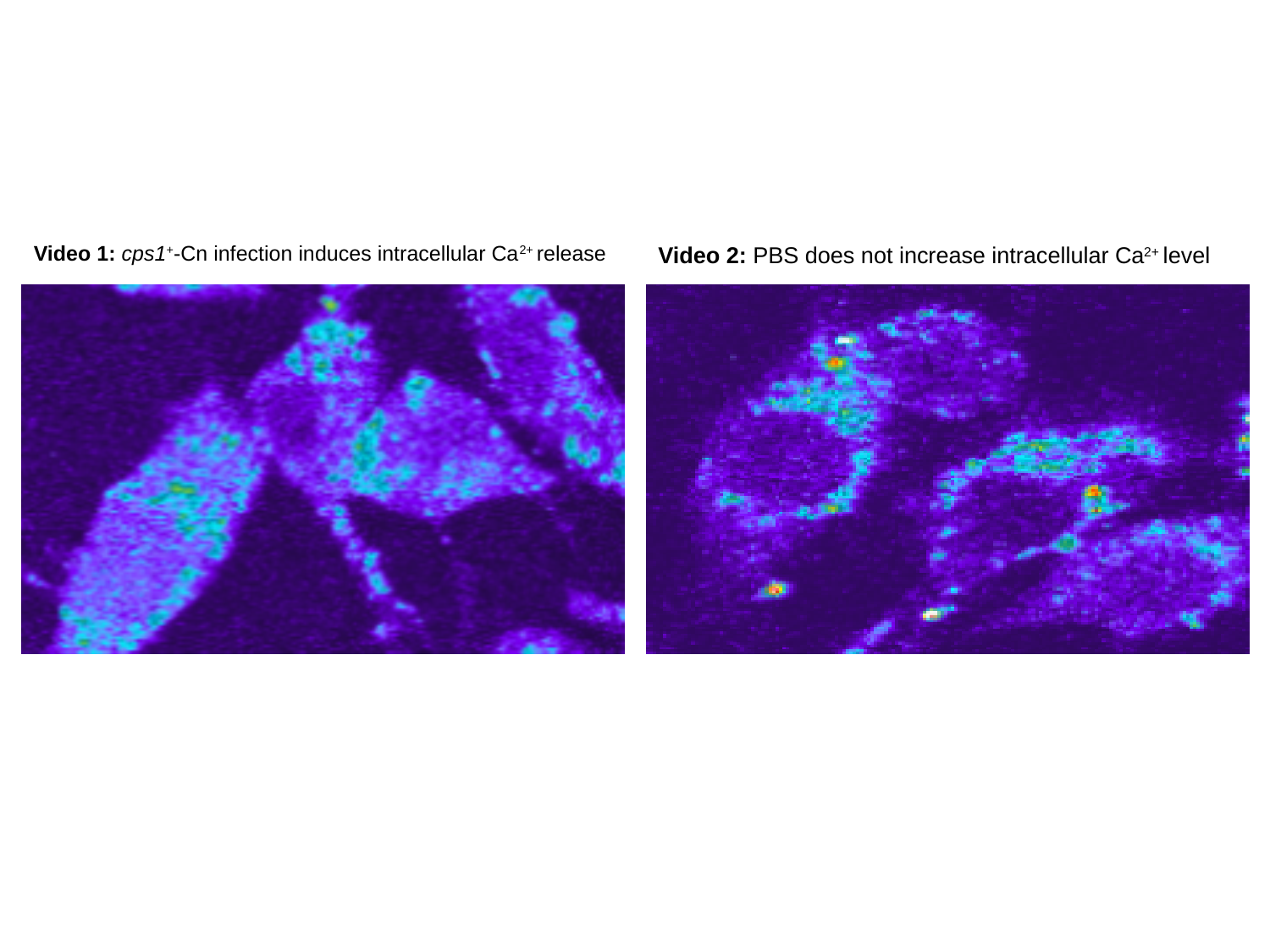

Video 1: cps1^+^-Cn infection induces intracellular Ca^2+^ release
Video 2: PBS does not increase intracellular Ca^2+^ level
